# Leishmania guyanensis controls endogenous viral replication by a canonical RNA interference pathway

**DOI:** 10.64898/2026.08.09.743808

**Authors:** Donnamae Klocek, Rhys H. Parry, Grant A. Kay, Aditya Reddy, Edubiel A. Alpizar-Sosa, Kristína Záhonová, Aitor Casas-Sanchez, Jovana Sádlová, Petr Volf, Alain Kohl, Vyacheslav Yurchenko

## Abstract

Protistan parasites of the genus Leishmania, infamous human and animal pathogens, can themselves be infected by endosymbiotic viruses, exemplified by Leishmania RNA viruses (LRVs). These viruses affect immune responses in vertebrate hosts and have been associated with adverse treatment outcomes. How parasites control replication of these viruses is not known. Intriguingly, functional RNA interference (RNAi) pathways that have been associated with antiviral responses across eukaryotes, are retained only in some *Leishmania* spp., including those of the subgenus *Viannia*. Here, we investigated effectors in the canonical RNAi response and the Piwi protein of the human pathogen *L.* (*Viannia*) *guyanensis* by gene ablation and identified Dicer-like 1 and Argonaute 1 proteins of the canonical RNAi pathway as critical for controlling viral RNA levels. Notably, we characterized virus-derived small interfering RNA (vsiRNA) levels and their unique properties including terminal modifications as well as, unusual for canonical Dicer cleavage, predominant perfectly matching sequence overlaps in blunt ended vsiRNA duplexes. Taken together, the data suggests that control of viral replication is directly mediated by the canonical RNAi response. This study opens the door to further investigations of antiviral RNAi in other protistan parasites and suggests that, where present, canonical RNAi is critical for such activities.

**Author summary:** *Leishmania* parasites of humans and animals harbor endosymbiotic viruses, which, in some cases, have been shown to affect vertebrate immune responses and impact treatment. Thus, understanding how viral levels are controlled is critical to identify antiviral effectors, which, in turn, will allow studies on how viral levels impact parasite biology. Here, we investigated RNA interference pathways against its virus of the family *Pseudototiviridae* in a New World human pathogen *L. guyanensis*. To do that, we have produced and analyzed genetic knockouts of Dicer-like and Argonaute proteins involved in antiviral small RNA response.

**Graphical Abstract:** 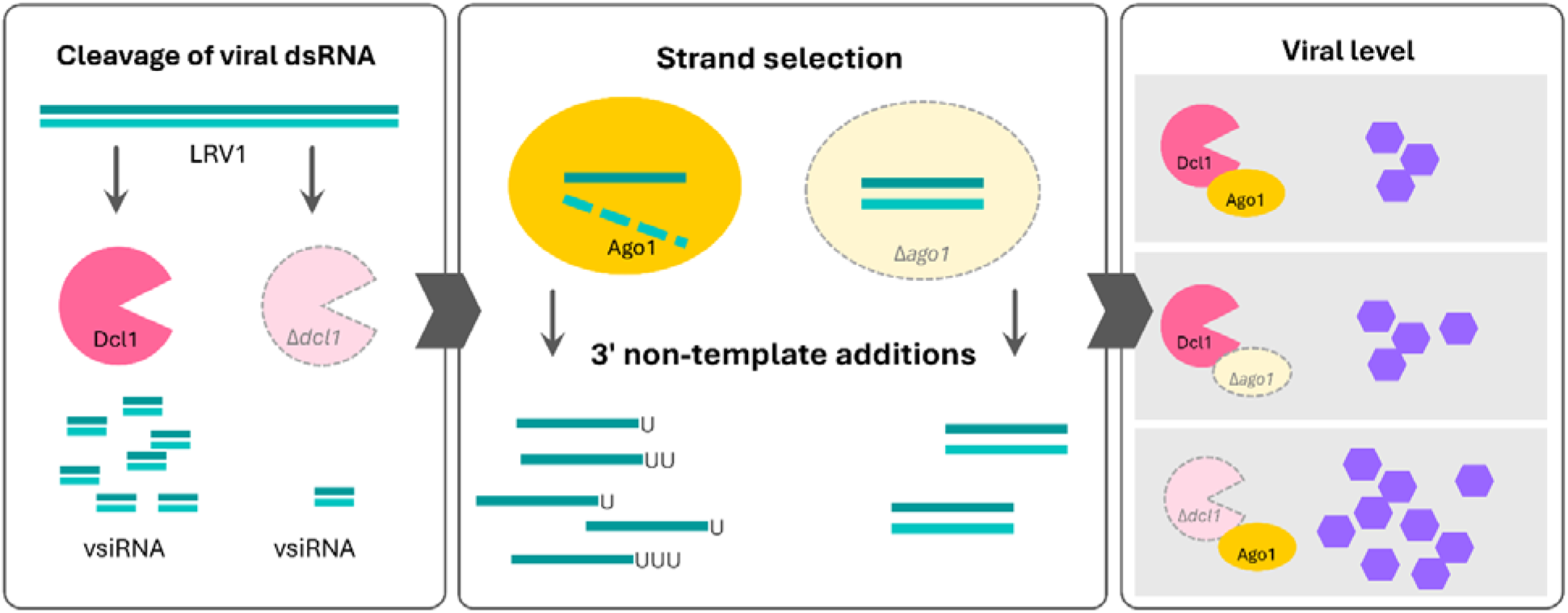

## Introduction

Parasites of the genus *Leishmania* (Euglenozoa: Trypanosomatidae [1]) cause cutaneous, mucocutaneous, and visceral leishmaniasis in vertebrates after, typically, sand fly transmission. They are mostly restricted to the tropical and subtropical regions and constitute a major public health issue [2–4]. Intriguingly, many, though not all, harbor viruses that replicate in the cytosol, such as *Leishmaniavirus ichi* (*Leishmania RNA virus 1*, LRV1), *Leishmaniavirus ni* and *Leishmaniavirus sani* (formerly united under *Leishmania RNA virus 2*, LRV2) of the family *Pseudototiviridae* with a dsRNA genome [5,6]. *Leishmaniavirus ichi* is restricted to species of the *Leishmania* subgenus *Viannia*, while *L. ni* and *L. sani* infect different species of the subgenus *L.* (*Leishmania*) [7]. Of note, two other leishmaniaviral species, which are phylogenetically close to *L. ni* and *L. sani*, were documented to infect parasites of the subgenus *L.* (*Sauroleishmania*) [8] and *Trypanosoma platydactyli* [9], respectively, while three more distantly related *Leishmaniavirus* spp. can infect flea-dwelling trypanosomatids of the genus *Blechomonas* [10]. The underlying biology and evolution of these endosymbiotic virus-parasite interactions as well as host control remain poorly understood and understudied [6,11–15]. The presence of LRVs has been shown to impact interaction with host cells *via* TLR (Toll-like receptor)-dependent pathways in vertebrates leading to exacerbated disease [16] as well as impacting treatment outcomes in some studies [17–19].

Notably, *Leishmania* subgenera harboring these viruses have either lost (*Leishmania* and *Sauroleishmania*) or retained (*Viannia*) canonical small interfering RNA-based RNA interference (RNAi) pathways involving Argonaute (Ago1) and two Dicer-like (Dcl1/Dcl2) proteins [20–23], a pattern also observed across other protistan parasites [24–27]. In *L. guyanensis* and *L. braziliensis*, where canonical RNAi proteins are present, virus-derived small interfering RNAs (vsiRNAs) from LRV1 have been detected, but the origin of these small RNAs as well as the mechanisms governing actual antiviral activity remain unclear. However, when the RNAi machinery is induced by dsRNA, it is capable of clearing the virus, which strongly suggests that the small RNA response is anti-virual [28]. Thus, the canonical RNAi in these species (initiated by Dcl cleavage of virus-derived dsRNA into vsiRNAs that are taken up by Argonaute to degrade matching target viral RNA) is comparable to antiviral RNAi across eukaryotes in general [29–31]. How species in the subgenus *Leishmania* deal with viral replication in the absence of a canonical RNAi machinery remains an open and intriguing question [32–34]. In addition, in iconic *T. brucei*, Dcl1 and Dcl2 were previously described as cytosolic and nuclear, respectively [35,36], suggesting distinct roles for these proteins as effectors [25]. Thus, there remain critical knowledge gaps in the functionality of RNAi effector proteins, in particular, concerning their role(s) in *Leishmania* antiviral defense.

Beyond canonical RNAi, the Piwi pathway, traditionally associated with genome stability and repression of transposable elements [37–39], is barely explored in kinetoplastids. It gives rise to a class of single stranded small RNAs (piRNAs) produced from double stranded RNA in ciliates, and linear RNA in animals [40]. Virus-derived piRNAs have also been described in insects, cnidarians, echinoderms, and mollusks, suggesting a wider adaptation of such a response to control viral infections [41,42]. They are generally from 21 nt to over 30 nt in length with a preference for U in position 1, and a so-called ping-pong amplification enriching A in position 10. There are, however, differences and in nematodes, where *Caenorhabditis elegans* piRNAs show a U1 bias only [43–46]. Ago and Piwi proteins of the piRNA pathway form subgroups within the PAZ/Piwi protein family, with the C-terminal Piwi domain resembling RNase H folds, which are important for slicer activity [47,48]. In the context of trypanosomatids, a Piwi protein (with a divergent PAZ domain) had been described in *T. cruzi* [49] and implicated in a pathway that has been suggested to retain aspects of siRNA and canonical piRNA pathways [50]. Similarly, the *T. brucei* genome encodes a Piwi protein that appears not to be involved in RNAi but instead controls mitosis and chromosome segregation [51]. Thus, much about the roles and functions of such proteins in trypanosomes remains to be discovered. Piwi proteins have also been documented in *L. major* and *L. infantum* (both from subgenus *Leishmania*), however, called Piwi-like proteins because of sequence and domain divergence. These proteins lack Ago and Dcl domains, retaining only a Piwi domain as a recognizable feature. Deletion of the Piwi-encoding gene results in viable parasites that display growth defects in amastigotes (possibly linked to dysregulation of genes involved in development), as well as delayed disease pathology in mice [52]. It is clear that many important questions on the roles and functions of Piwi proteins in *Leishmania* remain unanswered, including their involvement in antiviral responses.

Beyond the primary processing of small RNAs, post-transcriptional modifications, such as non-template additions (NTAs) of nt, may represent an additional layer of small RNA regulation. Indeed, ∼20% of *L. braziliensis* siRNAs already carry non-templated 3’ U due to the absence of HEN1 methyltransferase protection [22], suggesting that such NTAs could be a small RNA regulatory mechanism in *Leishmania* spp. that remains to be explored in the context of viral infection.

In this work, we investigated the role(s) of canonical RNAi and the Piwi protein in antiviral responses in new world *L. guyanensis*, which hosts *Leishmaniavirus ichi* LRV1-4. We produced genetic knock-outs (KOs) of RNAi components to assess the impact of this pathway on viral replication and parasite biology, implicating the canonical RNAi proteins Ago1 and Dc1 in an antiviral response against LRV. We also analyzed the nature and composition of small RNAs in these lines to further understand production and modifications of small RNAs during vsiRNA biogenesis.

## Results

### Identification of L. guyanensis RNAi effectors

To assess the components of RNAi machinery across Kinetoplastea species, we conducted genome wide searches for Ago1, Dcl1 and Dcl2, Rif4/5 [24], as well as Piwi genes. This identified presence, absence, or pseudogenization across a number of species (Fig 1, S2 Table).

**Fig 1.**
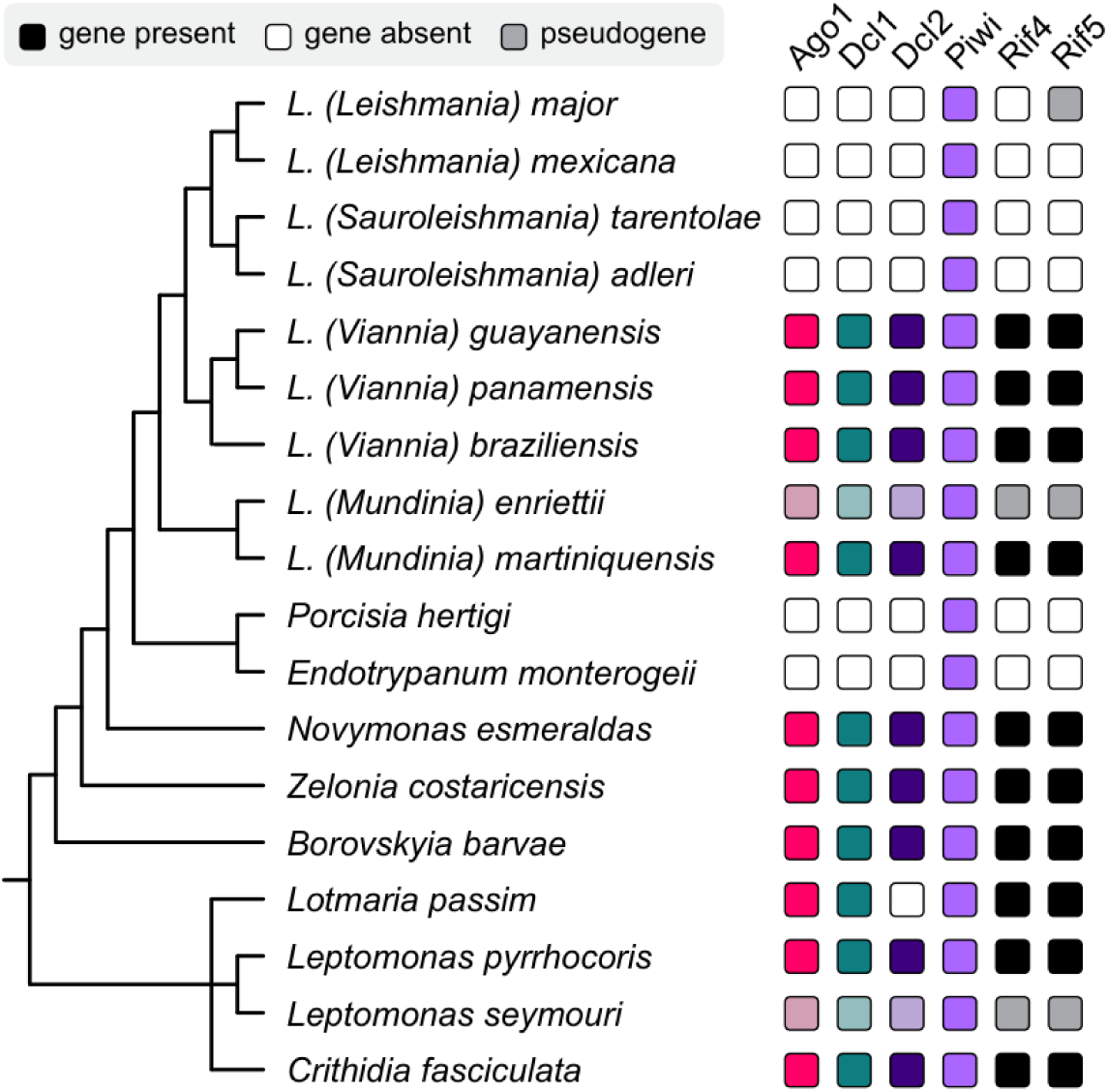
RNAi effectors in the Leishmaniinae. Identified and missing proteins, as well as pseudogenes are shown as black, white, and grey squares as explained in the graphical legend. Species relationships are shown on left by a schematic phylogenetic tree based on previous work [53].

The presence of the same protein domains across *L. guyanensis* and *T. brucei* RNAi proteins (Fig 2) indicates that the core components of the RNA targeting machinery are conserved across distantly related species, suggesting functional conservation of proteins. Supporting this, RNAi proteins generally cluster by species relatedness (S1-S4 Figs).

**Fig 2.**
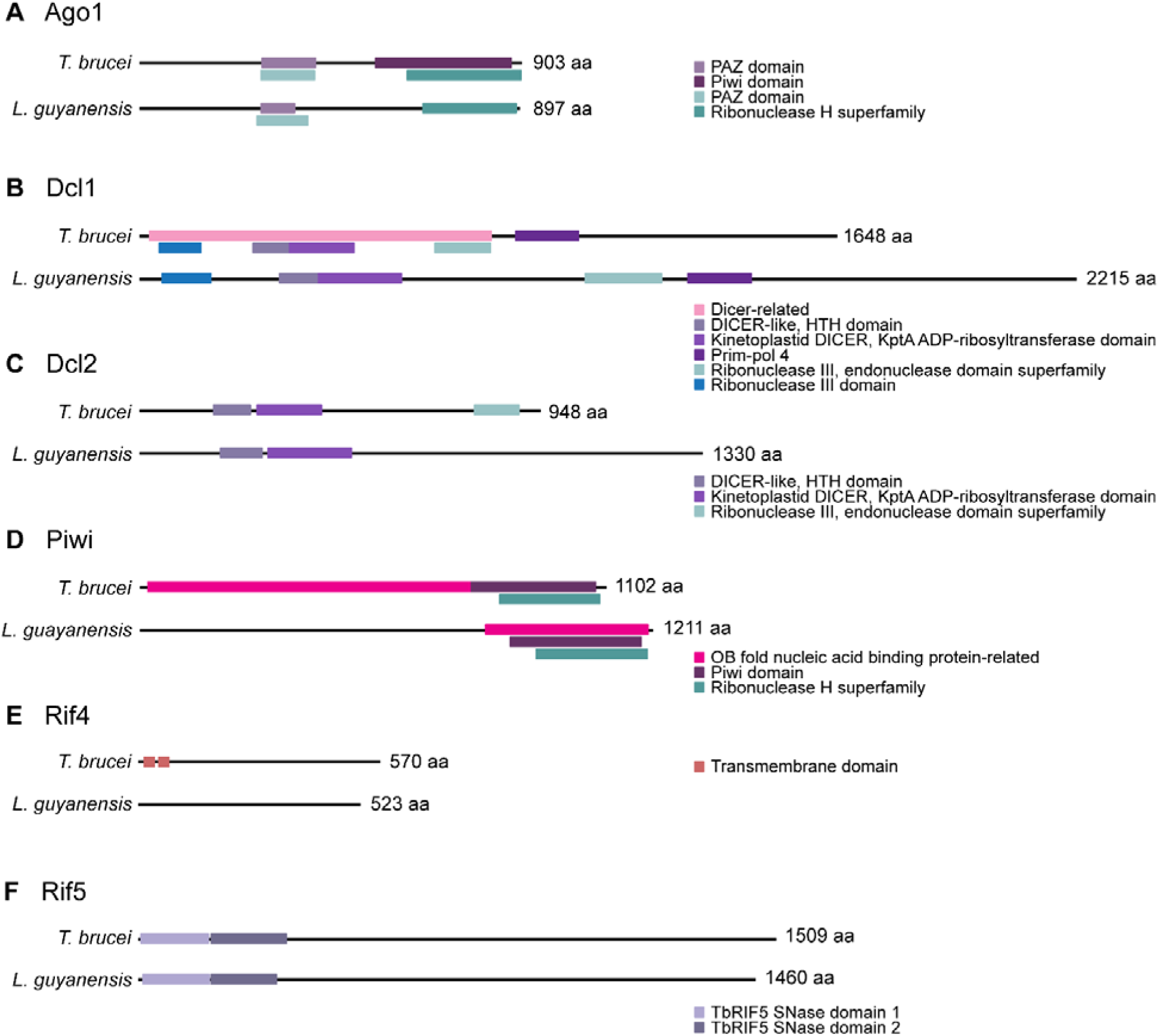
Predicted protein domains of RNAi pathway effectors Ago1, Dcl1, Dcl2, Piwi, Rif4, and Rif5. Shown are comparisons of domain structures of *T. brucei* and *L. guyanensis*. Domains in pink, purple, and cyan shades were predicted by Panther, Pfam, and Gene3D, respectively. Prediction of the ribonuclease III domain in Dcl1 (blue) is based on previously published work [36].

*Ablation of canonical RNAi and Piwi genes from L. guyanensis produces viable mutants* Deletions were introduced into both alleles of *L. guyanensis dcl1*, *dcl2*, *ago1*, and *piwi* genes to generate KO lines using a Cas9-based system adapted for trypanosomatids, yielding viable mutants and confirming that each of these genes is non-essential, at least in culture. Deletions were confirmed by genome sequencing (S5 Fig).

The *Δdcl1* and *Δdcl2* lines exhibited division rates comparable to or slightly higher than that observed in the parental WT line culture under similar conditions. Conversely, the *Δago1* and *Δpiwi* lines showed slower growth rates relative to the parental line, with the defect being particularly pronounced in the *Δago* line (S6A Fig). *Δdcl2* parasites showed morphological measurements comparable to those of the parental WT line, with the exception of slightly longer cell bodies and a corresponding increase in the nucleus–anterior distance. Both *Δdcl1* and *Δago1* lines had shorter but wider cell bodies, shorter flagella, and smaller nuclei than the parental line, with the anomalies being more prominent in the *Δago1* line. *Δpiwi* parasites displayed the most pronounced morphological changes, with all morphometric values except kinetoplast position reduced relative to the parental line (S6B Fig).

### RNA level is increased upon dcl1 and ago1 deletion

As with comparable viruses, such as the closely related totivirus ScL-A in yeast *Saccharomyces cerevisiae*, LRV1-4 exists in a dynamic tension with its *L. guyanensis* host, its real-time replication rate shifting with the ratios of, and complex interplay between, proviral and antiviral host factors that enable or limit its replication, respectively [54–56]. Owing to natural stochasticity in host gene expression, even minor early differences in the pro- versus antiviral factor balance can become amplified over time, producing substantial variation in viral levels [57]. An antiviral response through RNAi would target viral RNAs; as such, ablation of these genes should result in an increase in viral levels above baseline variance. LRV1-4 level was measured in three subcultures of each of the WT and KO lines at log phase by quantitative RT-PCR of viral RdRP to quantify viral load (Fig 3). The Δ*dcl1* line exhibited a four-fold higher level of LRV1-4 than the parental WT line, indicating that control of viral replication was strongly reduced in this KO line. The Δ*dcl2* line did not show any significant change. This reciprocal effect of the *dcl* mutations on LRV1-4 level mirrors that observed with corresponding mutations in *T. brucei*, where ablation of *Tb*Dcl1 rendered cells unresponsive to challenge with exogenous dsRNA, whereas ablation of *Tb*Dcl2 potentiated dsRNA-triggered RNAi activity [35]. Notably, LRV1-4 level in the Δ*ago1* line was also significantly higher compared to the parental control. This is consistent with a critical antiviral role for canonical RNAi, with Dcl1 being a key for vsiRNA production, and Ago1 acting downstream as vsiRNA-guided effector targeting viral RNA for degradation [30,58].

**Fig 3.**
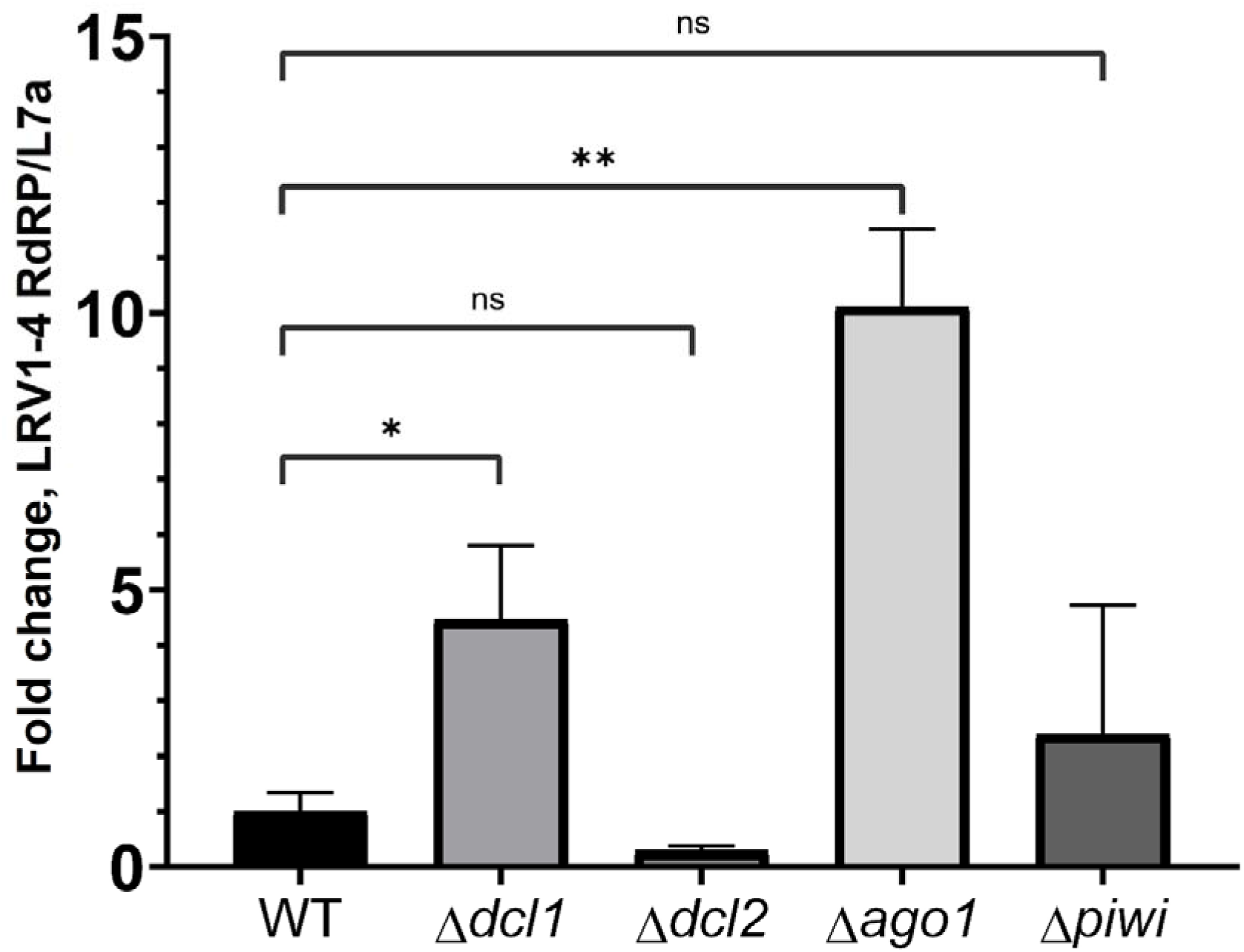
Viral load analysis. Viral levels in the WT and KO lines were normalized to expression of the ribosomal protein L7a gene. Statistical analyses were done using the Welch’s *t*-tests (*p*-values are as follows: *, < 0.05; **, < 0.01; ns, non-significant). Error bars represent standard deviations (SD).

### Roles of RNAi effectors and KOs on LRV1-4 vsiRNAs

The impact of RNAi effectors on small RNA, and LRV1-4 derived small RNA production was investigated using high-throughput small RNA sequencing from the WT parental and KO lines of *L. guyanensis*. For greater depth, two clones of each KO line were used and compiled for all RNA analyses. Raw populations of small RNA read lengths were between 15-30 nt that are prototypical for *Leishmania* spp. (S7 Fig). The WT libraries displayed characteristic 22 nt peaks with strong 5′ U bias, while *ago1* and *dcl1* deletions showed dramatic shifts to shorter size classes with peaks at 15-18 nt, accompanied by altered nucleotide bias toward G and gradual tailing off toward longer sizes.

However, *dcl2* deletion maintained more canonical size ranges with preserved U bias, while *piwi* deletion closely resembled WT profiles. Examination of perfectly mapped siRNA size classes in WT and RNAi effector KO lines (Fig 4A) indicated that LRV-derived small RNAs in libraries showed characteristic 21-23 nt vsiRNA peaks, consistent with previous conservations for LRV1 [28]. In the WT libraries forward-mapping reads show characteristic 3′ U bias, while reverse-mapping reads display 3′ A bias. Coverage analysis of mapped 21-23 nt reads (Fig 4B) revealed relatively uniform vsiRNA distribution across the LRV1-4 genome with both sense and antisense strands targeted, supporting bidirectional dsRNA substrate recognition by the antiviral machinery.

**Fig 4.**
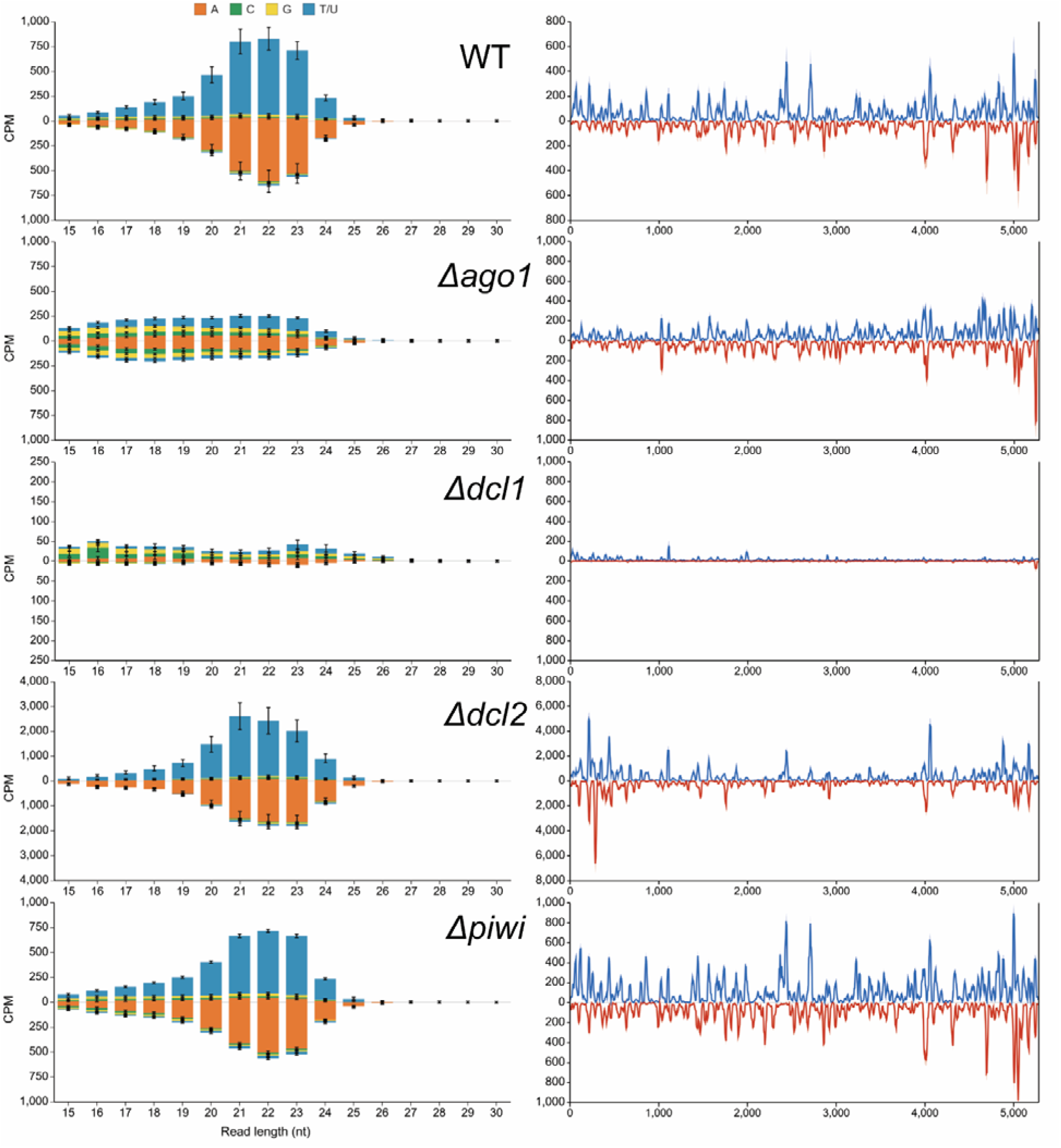
RNAi effector KOs reveal distinct roles in LRV1-4-derived vsiRNA processing and 5′ nt signatures. **(A)** Small RNA sequencing of *L. guyanensis* WT and RNAi effector KO lines, as indicated, that are infected with LRV1-4. Histogram of perfectly mapped small RNA reads 15-30 nt in length, mapped to the LRV1-4 genome strand (NC_003601.1, positive y-axis) and reverse strand (negative y-axis), with colors indicating 5′ terminal nucleotide prevalence per size class. Data shown as mean counts per million (CPM) of mapped reads from WT controls (n=4, ±SD) and KO lines generated from two independent clones with four biological replicates each (n=8 per KO). **(B)** Coverage profiles of 21-23 nt vsiRNAs mapped across the LRV1-4 genome (blue, sense) and antigenome (red, antisense), showing genome-wide distribution patterns and relative abundance differences between genotypes (y-axis, normalized read coverage ± SD). Y axis scales for CPM differ between lines.

Consistent with these size-distribution patterns, quantitative analysis of total 21–23 nt LRV1- 4-derived vsiRNA abundance showed that *dcl1* loss had the strongest effect, reducing canonical vsiRNAs by approximately 20-fold relative to WT (*p* < 0.0001, ordinary one-way ANOVA with uncorrected Fisher’s LSD; S8 Fig), while completely disrupting 5′ nt bias patterns (Fig 4A). Residual reads shifted toward 23 nt, suggesting Dcl2 attempts to compensate for loss of Dcl1 activity, but with altered cleavage specificity. Conversely, the absence of Dcl2 resulted in significantly increased vsiRNA abundance (∼5-fold, *p* < 0.0001; S8 Fig) with a weak shift toward 21 nt reads. The absence of Ago1 produced a more moderate phenotype: significantly reduced vsiRNA abundance (*p* < 0.05; S8 Fig) distributed across a broader, more diffuse 17-23 nt size range with diminished 5′ nucleotide bias. The absence of Piwi showed no discernible impact on vsiRNA production, size distribution, or nucleotide bias (*p* > 0.05), further excluding this protein from antiviral RNAi activity against LRV1-4 (Fig 4 and S8 Fig).

### Contribution of RNAi effectors to viral derived sRNA 3′ non-template additions in L. guyanensis

Non-templated 3′ uridylation has previously been reported in leishmanial small RNAs, with approximately 20% of *L. braziliensis* siRNAs carrying 3′ non-templated U [22]. To test whether a similar pattern occurs in LRV1-4-derived vsiRNAs in *L. guyanensis* and whether it depends on RNAi pathway proteins, we examined the frequency and composition of 3′ non-template additions (NTAs) across WT control and KO lines (Fig 5A). Most LRV1-4-derived vsiRNAs mapped perfectly, while a smaller, but reproducible, fraction carried 3′ NTAs. In WT controls, 87.7% of vsiRNA reads mapped perfectly and 12.3% contained a 3′ extension. In WT line, NTAs were predominantly mononucleotide additions, with much smaller contributions from di-, tri-, and tetranucleotide tails. Most LRV1-4- derived vsiRNAs mapped perfectly, while a smaller but reproducible fraction carried 3′ NTAs. In WT controls, 87.7% of vsiRNA reads mapped perfectly and 12.3% contained a 3′ extension. Similar perfect/NTA proportions were observed across KO lines; however, these proportions do not reflect total vsiRNA abundance, which was strongly reduced in *Δdcl1* (S8 Fig). Among NTA-containing reads, WT, *Δago1*, *Δdcl2*, and *Δpiwi* libraries were dominated by mononucleotide additions, whereas the residual Δdcl1 NTA-positive reads showed a shift towards longer tail classes, particularly tetranucleotide additions (Fig 5A).

**Fig 5.**
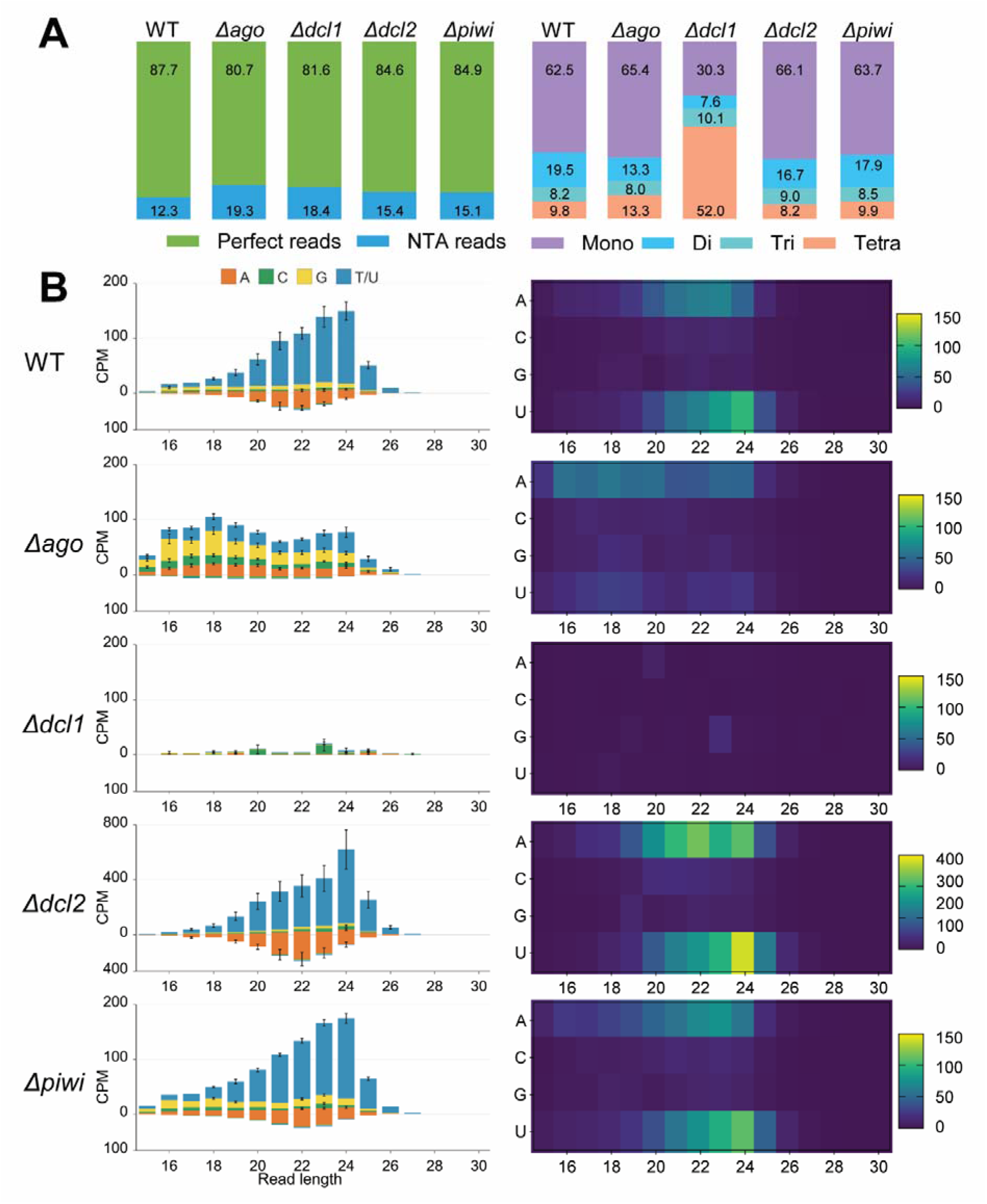
RNAi effector proteins shape the abundance and composition of 3′ NTAs in LRV1-4-derived vsiRNAs. **(A)** Proportion of perfectly mapped versus 3′ NTA-containing LRV1-4-vsiRNA reads in WT control and RNAi KO lines (left), and distribution of NTA classes by tail length as mono-, di-, tri-, or tetranucleotide additions (right). **(B)** Read-length distributions of reads with NTAs (soft-clipped reads) LRV1-4-derived vsiRNAs carrying mononucleotide 3′ NTAs only (left) and heat maps of the identity of the added terminal nt across read lengths, normalized as mean CPM (right). Stacked bars indicate CPM values ± SD for each size class colored by 5′ nucleotide (A, C, G, or U) bias. Data are shown as means for WT controls (n = 4) and KO lines (as indicated) generated from two independent clones with four biological replicates (n = 8).

To examine the composition of tailed vsiRNAs specifically, we analyzed only reads with NTAs (soft-clipped reads) carrying mononucleotide 3′ NTAs (Fig 5B). They displayed a distinct size profile, with the enrichment of longer species dominated by positive-sense vsiRNAs peaking at 23–24 nt.

Negative-sense vsiRNAs carrying 3′ NTAs were less abundant overall and showed a similar length bias to non-NTA reads (Fig 4), without a clear size-specific enrichment. The identity of the added nt was strongly biased towards U and to a lesser extent, A in WT controls, particularly among 23–24 nt reads, indicating that 3′ uridylation and adenylation are the predominant terminal modifications of LRV1-4-derived vsiRNAs in WT parasites. The loss of Ago1 abolished this clear mononucleotide and dinucleotide U-bias NTA signature (S9 Fig) despite a slight increase of the proportions of reads with NTA (19.3%), indicating that Ago1 is not simply required for vsiRNA production but, instead, is important for the accumulation of uridylated vsiRNAs. The *Δdcl1* line produced very few vsiRNAs overall and, correspondingly, little detectable NTA signal. Together, these data show that 3′ uridylation is a regular feature of LRV1-4 vsiRNA metabolism in *L. guyanensis* and that Ago1 makes a major contribution to this modification pathway.

### Ago1 deletion reveals accumulation of complementary vsiRNA overlap signatures

Overlap signature analysis of perfectly mapped vsiRNAs revealed genotype-dependent differences in complementary vsiRNA pair accumulation in *L. guyanensis* (Fig 6). WT libraries displayed strong depletion of perfectly overlapping vsiRNA pairs across most size classes, indicating active duplex unwinding and guide strand selection. Notably, the overlapping pairs shift of n-2 for small RNA read sizes (for example, a 21 nt siRNA duplex overlapping by 19 nt) seen in insects and other species [31,59–61] was not a dominant feature of LRV1-4-derived vsiRNAs, suggesting distinct dsRNA processing mechanisms. In contrast, *Δago1* libraries showed a clear enrichment of complementary overlap signals, centred around near full-length/blunt-ended overlaps but with some adjacent size heterogeneity. This pattern is consistent with accumulation of unresolved vsiRNA duplex-like species in the absence of Ago1, rather than complete loss of vsiRNA production. The fuzziness around the main enriched overlap signal likely reflects heterogeneity in vsiRNA size, terminal processing, or partial trimming of duplex-associated reads. Together, these data support a model in which Dcl1- dependent vsiRNAs are generated as complementary sense–antisense species, while Ago1 acts downstream to resolve these duplexes and select guide strands for antiviral RNAi activity.

**Fig 6.**
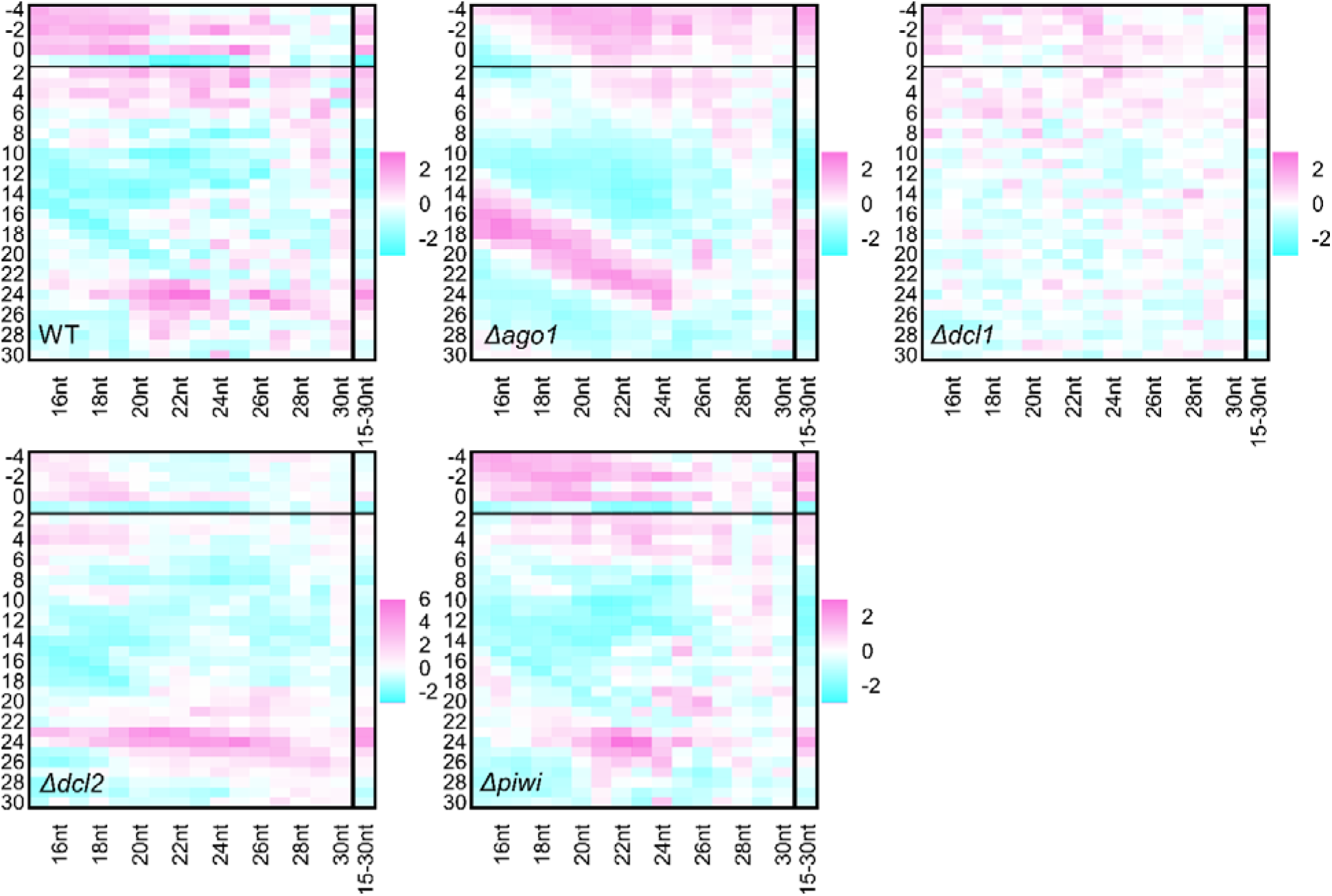
Deletion of *ago1* reveals intact vsiRNA duplexes through overlap signature analysis. Heatmaps showing overlap probability z-scores for perfectly mapped vsiRNAs (15-30 nt) across RNAi effector KOs. Purple indicates positive z-scores (enriched overlaps), blue indicates negative z-scores (depleted overlaps). Data represent mean z-scores from two independent clones with four biological replicates each (n = 8 per KO, n = 4 for WT).

Size-specific processing was also evident, with vsiRNAs ≥ 23 nt showing altered overlap signatures including slight enrichment for 24 nt overlaps, potentially indicating alternative processing pathways or increased duplex stability for longer vsiRNA species.

### Transcriptome-wide differential expression analysis across RNAi pathway KOs

To examine the host transcriptional responses to the RNAi effector ablations, we performed RNA sequencing. Library quality and sequencing depth were consistent across all samples and treatments (Fig 7A), with post-normalization distributions uniform across all libraries (Fig 7B). Multidimensional scaling of the normalized expression data revealed discrete separation of transcriptional response for all four KOs from WT along the first principal dimension (37.4% of leading log₂FC variance), with negligible dimension one within-group variation of the MDS, validating the experimental replicates (Fig 7C). Pairwise differential expression analysis of the KO treatments against WT (FDR < 0.05, log₂FC > 1) revealed subtle, but reproducible, differentially expressed genes responses across the four KOs (Fig 7D). The *Δago1* mutant showed the largest response with 74 DEGs (differentially expressed genes) (53 upregulated, 21 downregulated), followed by *Δpiwi* with 45 DEGs (15 upregulated, 30 downregulated) and *Δdcl1* with 42 DEGs (33 upregulated, 9 downregulated) lines. The *Δdcl2* mutant had the smallest number of differentially expressed genes compared to WT with only 10 DEGs (3 upregulated, 7 downregulated).

**Fig 7.**
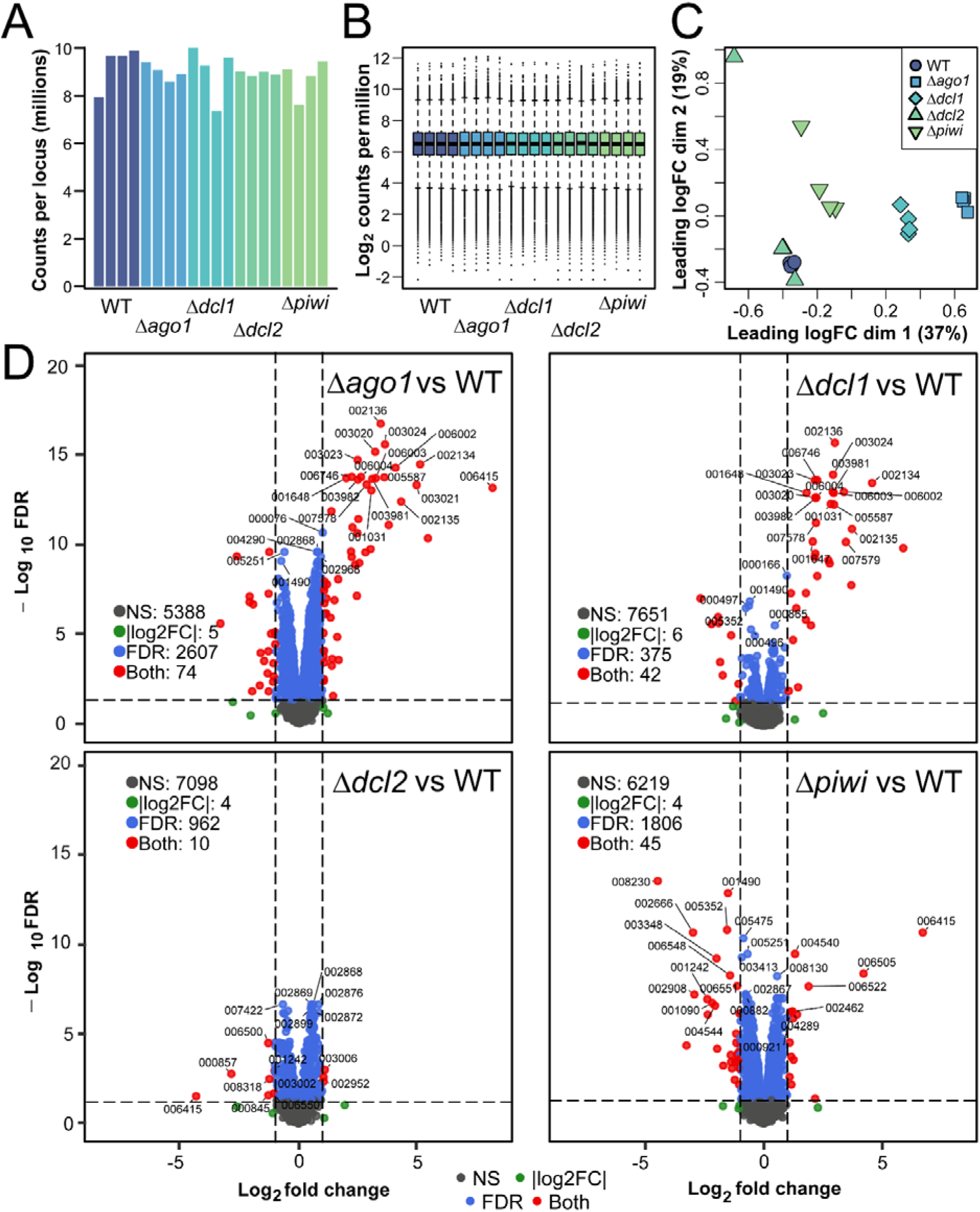
Transcriptome-wide differential expression analysis of RNAi pathway KO lines in *L. guyanensis*. **(A)** Total counts per locus across WT, *Δago1*, *Δdcl1*, *Δdcl2*, and *Δpiwi* RNA sequencing libraries. **(B)** Distribution of normalized gene expression values shown as log2 CPM for all libraries. **(C)** Multidimensional scaling (MDS) plot of normalized expression profiles. Percentages indicate the proportion of leading log_2_ fold change variance explained by each dimension. **(D)** Volcano plots showing pairwise differential gene expression for *Δago1*, *Δdcl1*, *Δdcl2*, and *Δpiwi* relative to WT. Dashed vertical lines indicate log_2_ fold change cut-offs and the dashed horizontal line indicates the FDR threshold. Points are colored according to significance category: not significant (NS), log_2_ fold change (log_2_FC) only, FDR only, or both thresholds. For readability, only representative high-confidence DEGs are labelled by locus suffix; complete pairwise differential expression results are provided in S6–S9 Tables.

### Shared transcriptional signatures in *Δago1* and *Δdcl1 L. guyanensis* are associated with transposable element de-repression

To assess how similar up and downregulated genes are between KO mutants, we performed a comparison of their gene sets using Venn analysis (S10A Fig). We identified that DEG profiles of *Δago1* and *Δdcl1* are similar with 32 genes commonly upregulated and six commonly downregulated in both KOs, with 21 genes uniquely upregulated and 15 uniquely downregulated in *Δago1*, and only one uniquely upregulated and three uniquely downregulated in *Δdcl1* (S6-S9 Tables). The log₂FC values of all 38 shared DEGs were strikingly correlated between the two comparisons (S10B Fig), with conserved upregulated genes spanning log₂FC 1.16–8.16 in *Δago1* and 1.07–5.91 in *Δdcl1*. The conserved upregulated loci were overwhelmingly composed of TATE retrotransposon components, one of the dominant mobile element families in *Leishmania* [62] accounting for 13 of the 32 shared upregulated genes, including TATE gag proteins (NXY56_002134, NXY56_006002), reverse transcriptases (NXY56_003021, NXY56_003024, NXY56_006004, NXY56_006006, NXY56_007578, NXY56_001648, NXY56_005588), tyrosine recombinases (NXY56_006003, NXY56_003020, NXY56_003023, NXY56_000167), and a phage integrase family protein (NXY56_000779), consistent with RNAi-mediated suppression of transposable elements as a primary pathway function. Additional TATE gag and reverse transcriptase genes were present among *Δago1*-unique upregulated genes (NXY56_003022, NXY56_003025, NXY56_001030). Taken together, these results support a model, in which Dcl1 and Ago1 act as core components of a shared RNAi pathway that also mediates repression of transposable element transcripts in *L. guyanensis*.

## Discussion

In this study, we investigated the antiviral roles of the RNAi pathway against LRV1-4 in *L. guyanensis*. Genetic ablations of the canonical RNAi proteins Dcl1, Dcl2 and Ago1, as well as of Piwi, were generated. Critically, the *Δdcl1* and *Δago1* lines showed significantly increased viral RNA levels. This was further confirmed by the vsiRNA analysis documenting read mapping to the full lengths of viral genome, implying that viral dsRNA must be accessible (likely, at some point during replication) to the RNAi machinery instead of being completely protected by the capsid. This could be due to the capsid disintegration, fault structures, etc. [63]. The exact reason remains speculative but, importantly, our data establish Dcl1 as a key regulator? in vsiRNA production. Should these Dcl1 and Dcl2 resemble their counterparts in *T. brucei*, where Dcl1 is cytosolic and Dcl2 nuclear [35], a role for Dcl1 in targeting a virus replicating in the cytosol would not be surprising. Notably, deletion of Dcl2 leads to an increase of vsiRNAs, but not an increase in antiviral activity, which suggests that saturation, or maximum efficiency of the antiviral RNAi response have been reached. Why more vsiRNAs were observed following Dcl2 deletion remains unclear. It may be that an activity or function of Dcl2 inhibits Dcl1, for example by limiting specific interaction partner pools that are required for activity. Supporting this, more overlapping vsiRNAs are observed in the absence of Dcl2 (and the overlap pattern resembles, even though, not precisely, that of the Ago1 deletion), suggesting that Ago1 cannot take up more of these excess vsiRNAs for processing. The observed pattern of vsiRNA overlap evident in the Δ*ago1*, where passenger strand cleavage in the vsiRNA duplex no longer occurs and duplexes are expected to accumulate, suggests perfectly matching vsiRNA duplexes scenario rather than the presence of 3’ nt overhangs. This is unexpected for RNAi proteins [64] and remains to be investigated further.

Although small RNAs derived from LRV1 have been previously documented in *L. guyanensis* [28], our study defines that RNAi, with no obvious Piwi contribution, actually acts antivirally in a natural infection. Moreover, we show that mono-3′ uridylation and, to a lesser extent, mono- 3′adenylation are present in a small percentage of vsiRNAs, reminiscent of uridylation of small RNAs in *L. braziliensis* [22]. Why the deletion of Dcl1 involves a shift from mono- to tri-NTAs is not clear.

Equally, the exact effector enzymes mediating NTAs, and the potential consequences of NTAs and related shifts in vsiRNA length, warrant further investigation. However, our data demonstrate that the majority of vsiRNAs retain original length and NTA may just play a role in the natural turnover of these molecules [65]. How and why Dcl1 and Ago1 impact expression of TATE retrotransposon also remains to be investigated further. It may be explained by Dcl2 losing its function or, perhaps, Dcl1 playing a different role. However, ablation of Ago1 also led to the accumulation of transcripts and reduction of siRNAs for TATE and other mobile elements in *L. braziliensis* [22], suggesting a more general regulatory effect of RNAi on such genomics elements.

In summary, in this work, we define the canonical RNAi pathway and specifically the Dcl1- initiated arm as an antiviral response against a member of the *Pseudototiviridae, Leishmaniavirus ichi* (LRV1-4), in *Leishmania guyanensis*. We have also generated parasite lines with increased viral titers, which may be useful for future studies in vectors and vertebrate hosts to assess the potential roles of viruses beyond the parasite cells themselves. It needs to be stressed that establishment of viable *Leishmania* lines with increased viral load suggests that cellular response mechanisms resulting in elimination of kinetoplastid cells infected by the virus either never existed, have been lost in these flagellates, or that other (not yet identified) antiviral response mechanisms may be limiting any negative effects of viral replication. Indeed, *Leishmania* that have lost RNAi can be infected with viruses [66,67] and, presumably, mount antiviral responses that are yet to be identified. It needs to be seen to what extent the findings of this study apply to other viruses infecting trypanosomatids [7], but our work opens the door to investigating such questions.

## Materials and methods

### Parasite cultures and establishment of KO lines

A previously characterized LRV1-4-positive *L. guyanensis* (MHOM/BR/75/M4147) cell line episomally expressing Cas9 and T7 polymerase [68] was cultured in 9.9 g/l M199 (MilliporeSigma/ Merck, Burlington, USA) supplemented with 2 μg/ml hemin (Jena Bioscience, Jena, Germany), 10% heat-inactivated fetal bovine serum (FBS, BioSera Europe, Nuaillé, France), 2 μg/ml biopterin, 100 units/ml of penicillin, 100 μg/ml of streptomycin (all Thermo Fisher Scientific, Carlsbad, USA), and 50 µg/ml hygromycin B (Roche Life Science, Penzberg, Germany) at 26°C. The identity of the species was confirmed by amplification and sequencing of its 18S rRNA and gGAPDH genes using primers and conditions described previously [69].

Cas9-mediated deletions within the genes of interest – *dcl1*, *dcl2*, *ago1*, and *piwi* to make *Δago1*, *Δdcl1*, *Δdcl2*, and *Δpiwi* lines – were done as previously described [70]. In brief, for each gene of interest, *L. guyanensis* was transfected with PCR-generated templates for two guide RNAs and donor DNA containing the puromycin resistance gene. Gene-edited populations (knockout [KO] lines) were selected in liquid M199 supplemented with 100 µg/mL of puromycin (InvivoGen, San Diego, USA). Knockouts were confirmed *via* diagnostic PCR targeting the deleted regions and next-generation sequencing (NGS) as described previously [71]. Primers used in this paper are listed in the S1 Table. Mutants were cloned *via* serial dilution in liquid supplemented M199, as described previously [10,72].

### Morphometry and growth kinetics

For each parasite line, 6 measurements of 200 cells were taken as previously described [73]: cell body length and width, flagellum length, nuclear diameter, nuclear position, and kinetoplast position. For analysis of growth kinetics, parasites were seeded in triplicate at 5× 10^5^ cells per ml. Parasites were counted every 24 hours for 7 days using a hemocytometer as previously described [74].

### Quantification of viral RNA levels

As viral replication is stochastic (i.e., individual host cells infected by a virus can exhibit divergent frequencies and timings of viral replication surges) clonal cultures were used to minimize host heterogeneity, thereby allowing stochastic viral dynamics to be assessed independently. Clonal cultures of the 5 lines (wild-type (WT) and 4 KO lines) were split into 3 flasks at a concentration of 5× 10^5^ parasites per ml and incubated in parallel at 26°C. After 96 hours (late log phase), total RNA from 5× 10^7^ cells per flask were extracted with TRItidy G (Avantor, Radnor, USA), treated with Turbo DNase (Thermo Fisher Scientific), then converted to cDNA using random hexamer primers and RevertAid First Strand cDNA Synthesis Kit (Thermo Fisher Scientific) following the manufacturers’ protocols. Levels of LRV1-4 were measured *via* RT-qPCR using primers against the viral RNA-dependent RNA polymerase (RdRP) (S1 Table). Viral levels were normalized to the expression of the gene encoding ribosomal protein L7a [75] and are expressed as fold change relative to WT.

### Homology searches and phylogenetic analyses

Components of the RNAi pathway were identified by BLAST v. 2.8.1+ searches [76] using *T. brucei* proteins as queries and genomes, genome-derived proteomes, and transcriptomes of other kinetoplastids from NCBI, TriTrypDB release 68 [77], and previous publications [23,53,78,79] as databases (S2 Table). Sequences that were identified with an *e*-value threshold of 0.0005 were used to search the genome-derived proteome of *T. brucei* to confirm the orthology with an *e*-value threshold of 0.05. Either standalone BLAST or BLAST implemented in the AMOEBAE workflow [80] was used. Protein domains were predicted by InterProScan v. 5.77 [81] implemented in Geneious Prime v. 2025.21 [82]. Identified kinetoplastid sequences were aligned together with *T. brucei* proteins by MAFFT v. 7.458 [83] using the L-INS-i algorithm and ambiguously aligned positions were removed by trimAl v. 1.4 [84] using -gt 0.8 option. Maximum-likelihood phylogenetic analyses were performed in IQ-TREE v. 3.0.1 [85] using the PMSF method [86] with the LG+C60+G model, the guide tree inferred with the LG+G model, 1,000 replicates for ultrafast bootstraps [87] and Shimodaira– Hasegawa approximate likelihood ratio test (SH-aLRT) [88], and a maximum of 5,000 iterations.

### Next generation sequencing and analysis of genomic DNA

Genomic DNA from 1× 10^8^ cells was extracted from each KO line using DNeasy Blood & Tissue Kit (Qiagen, Hilden, Germany) and sequenced at Macrogen (Amsterdam, Netherlands) as previously described [89]. Reference genome of *L. guyanensis* strain MHOM/BR/75/M4147 was obtained from our previous assembly BioProject PRJNA808737 [68,90]. Analysis was performed as previously described [91,92] with modifications. Briefly, reads were mapped onto the reference genome assembly using the Burrows-Wheeler Aligner BWA-MEM [93,94] with index files generated as needed. PCR duplicates were removed using the Genome Analysis Toolkit GATK v. 4.1.4.1 [95].

Reference FASTA indexing was performed in SAMtools v. 1.21 [96]. Paired-end FASTQ files for each sample were mapped, converted to BAM format, and sorted. Output files were indexed and manually inspected to confirm gene deletions using the Integrative Genome Viewer IGV v. 2.8.9 [97], where the KO loci were curated and visualized against the parental WT and against each other. Code and related scripts used for genome alignments, as well as corresponding raw read counts, genome mapping statistics, and sample metadata for all lines are provided in the S3 Table and the GitHub repository (https://github.com/edubielalpizar/L.guyanensis_M4147_KOs).

### RNA sequencing for KO line characterization and differential gene expression (DEG) analysis

For each of the WT and KO lines, total RNA from 1.3× 10^8^ cells were isolated using RNeasy Mini Kit (Qiagen) in quadruplicate and sequenced at Macrogen Europe (Amsterdam, the Netherlands).

Strand-specific mRNA libraries were prepared using the TruSeq Stranded mRNA LT Sample Prep Kit (Illumina, San Diego, USA) according to the manufacturer’s protocol and sequenced on the NovaSeq X platform (Illumina) to produce 150 bp paired-end reads. Library orientation was reverse-stranded.

Basecalled FASTQ files were processed with fastp v. 0.23.2 [98] to trim adapter sequences, remove low-quality reads, and discard unpaired and shorter than 75 nt reads. For host transcriptome analysis, filtered reads were mapped to the *L. guyanensis* strain MHOM/BR/75/M4147 reference genome (GenBank accession GCA_024970365) using HISAT2 v. 2.2.1 [99] with the parameters --rna-strandness RF --dta --passthrough. The resulting alignments were sorted by genomic position with SAMtools v. 1.18. Raw read numbers, genome mapping statistics, and metadata are provided in S4 Table. Gene-level read counts were generated using HTSeq v. 2.0.2 [100] with the annotation file GCA_024970365.1_ASM2497036v1_genomic.gtf, using union mode, reverse-stranded counting, a minimum alignment quality threshold of 10, exon as the feature type, and locus_tag as the gene identifier. Downstream analyses were conducted in R v. 4.5.2 within RStudio 2025.09.2, using packages including edgeR v. 4.8.2, ggplot2 v.4.0.2, ggrepel v. 0.9.8, data.table v. 1.18.2.1, and tidyr v. 1.3.2 [101–103]. Individual count files were merged into a single gene-by-sample count matrix, and differential gene expression analysis was conducted first by filtering out lowly expressed genes using a minimum total count threshold of 10 across all samples. After that, counts were normalized using the trimmed mean of M-values method. Tagwise dispersions were estimated using estimateDisp with robust settings, and differential expression was tested using quasi-likelihood negative binomial generalized log-linear models implemented in glmQLFit and glmQLFTest of the edgeR package [104]. Differentially expressed genes were identified on the basis of log_2_ fold change and false discovery rate (FDR)-adjusted *p-*values. Visualization of log_2_ fold change and FDR-adjusted *p-*values was performed using volcano plots generated with EnhancedVolcano v. 1.28.2 [105].

### Small RNA sequencing and analysis

Total RNA was extracted in quadruplicates from 6.5× 10^7^ cells of each line using Zymo Direct-Zol RNA Miniprep Plus Kit (Zymo Research, Irvine, USA), treated on-column with DNase I as per the manufacturer’s instructions, and quantified using a Qubit RNA BR kit (Thermo Fisher Scientific).

Subsequent quality control, library preparation, and sequencing reactions were carried out by Lexogen (Vienna, Austria). Sampled integrity was assessed on a Fragment Analyzer System (Agilent Technologies, Santa Clara, USA) and library preparation was performed on 400 ng input RNA using Lexogen’s miRVEL Discovery Library Prep Kit and 12nt Unique Dual Indices according to the manufacturer’s protocol. cDNA libraries were sequenced using the Illumina NovaSeq X platform (Illumina) with single end 90 nt reads.

Raw sequencing reads were quality-controlled and adapter-trimmed using fastp with the 3’ adapter sequence (5’-AACTGTAGGCACCATCAAT-3’). Only reads containing the adapters that were 16 nt in length or greater and without ambiguous nucleotides were kept for downstream analysis. Final trimmed libraries ranged between 2.85 and 15.2 million reads with an average of 10.5 million reads per library (S5 Table). Clean, trimmed reads were mapped onto the *L. guyanensis* M4147 LRV1-4 genome (GenBank accession NC_003601) using Bowtie2 v. 2.4.5 [106] with local alignment parameters optimized to preserve non-template additions: --local -L 15 -N 1 --score-min G,10,0.

Mapped BAM files were processed using tailclip_filter.py to separate perfectly mapping reads (noClip.bam) from reads containing soft-clipped sequences indicative of non-template additions (withClip.bam). Scripts used for small RNA analysis are available from the viral_sRNA_tools GitHub repository at: https://github.com/rhparry/viral_sRNA_tools. Mononucleotide and dinucleotide addition frequencies were enumerated and normalized to input library size. Overlapping virus-derived small RNA pairs and overlap probabilities (z-score) were calculated iteratively from the noClip BAM files using the small RNA signatures Python script [107]. Rather than restricting the analysis to perfectly size-matched duplexes, overlap enrichment was assessed across LRV1-4-derived vsiRNAs spanning 15–30 nt to capture size-asymmetric and non-canonical pairing patterns. Analysis was performed in two modes: (i) focal read-length analysis, in which each individual query size class was tested against the full 15–30 nt complementary small RNA population using parameters such as --minquery X --maxquery X --mintarget 15 --maxtarget 30 --minscope -4 --maxscope 30, where X represents each individual size class from 15 to 30 nt; and (ii) collective analysis across the full size range using --minquery 15 --maxquery 30 --mintarget 15 --maxtarget 30 --minscope -4 --maxscope 30 to assess overall overlapping signature patterns across all vsiRNA size classes simultaneously. Two clones were analyzed for each KO line (C1/C2) and one for parental WT (C1).

## Data availability

RNA and DNA sequencing reads used in this study have been deposited in the Sequence Read Archive (SRA) [108] in the NCBI database under the BioProject accession numbers PRJNA1445940 (small RNA and RNA sequencing) and PRJNA1458557 (genome sequencing); for the latter, BAM files were also archived in Zenodo (https://doi.org/10.5281/zenodo.19185930).

## Acknowledgements

We thank members of our laboratories for stimulating discussions. Computational resources (Czech side) were provided by the e-INFRA CZ project (ID: 90254), supported by the Ministry of Education, Youth and Sports of the Czech Republic. Computational resources (Australian side) were also supported by the University of Queensland Research Computing Centre, including access to the Bunya high-performance computing facility.

## Author contributions

Conceptualization: VY, AK; data curation: DK, RHP, GAK, EAA-S, KZ, ACS, JS; formal analysis: DK, RHP, GAK, AR, EAA-S, KZ, ACS, JS; Investigation: DK, RHP, GAK, EAA-S, KZ, ACS, JS; funding acquisition: RHP, PV, AK, VY; methodology: RHP, EAA-S, ACS; supervision: PV, AK, VY; visualization: DK, RHP, EAA-S, KZ, AK, VY; writing –original draft: DK, RHP, GAK, KZ, AK, VY; writing –review & editing –all authors.

## Conflict of interest statement

None declared.

## Funding

Grant Agency of the Czech Republic [26-20420S to VY and PV], EU’s Operational Program ‘Just Transition’ [CZ.10.03.01/00/22_003/0000003 LERCO to VY], Australian National Health and Medical Research Council Ideas Grant [2038097 to RHP], Australian Research Council DECRA Fellowship [DE260101042 to RHP], LSTM Director’s Catalyst Award [to GAK], LSTM startup support [to AK].

Funding for open access charge: EU’s Operational Program ‘Just Transition’ [CZ.10.03.01/00/22_003/0000003 LERCO].

## Data availability

The data underlying this article are available in the article and in its online supplementary material. RNA and DNA sequencing reads used in this study have been deposited in the Sequence Read Archive in the NCBI database under the BioProject accession numbers PRJNA1445940 (small RNA and RNA sequencing) and PRJNA1458557 (genome sequencing); for the latter, BAM files were also archived in Zenodo (https://doi.org/10.5281/zenodo.19185930). Code and related scripts used for genome alignments are provided in the GitHub repository https://github.com/edubielalpizar/L.guyanensis_M4147_KOs Scripts used for small RNA analysis are available from the viral_sRNA_tools GitHub repository https://github.com/rhparry/viral_sRNA_tools

## Supporting information captions

**S1 Table. Primers used in this study.**

**S2 Table. RNAi core component proteins across kinetoplastids.** Species, database and dataset are listed together with accession numbers as indicated.

**S3 Table. Sequencing read processing summary.** Raw and trimmed read counts and read retention percentages are shown for each sample. Full mapping statistics are available *via* GitHub repository at https://github.com/edubielalpizar/L.guyanensis_M4147_KOs.

**S4 Table. Total RNA sequencing library metadata and genome mapping summary.** Library-specific metrics for each biological replicate including total mapped reads to the *L. guyanensis* genome used for DEG analysis.

**S5 Table. Small RNA library metadata and mapping analysis.** Library-specific metrics for each biological replicate including total mapped reads (15-30 nt) to the LRV1-4 genome, perfectly mapped reads (no mismatches), and soft-clipped reads (containing potential non-template additions).

**S6 Table.** Pairwise differential expression analysis of *Δago1* relative to WT. Gene-level differential expression results including log2 fold change, statistical test results, and FDR-adjusted p-values.

**S7 Table.** Pairwise differential expression analysis of *Δdcl1* relative to WT. Gene-level differential expression results including log2 fold change, statistical test results, and FDR-adjusted p-values.

**S8 Table.** Pairwise differential expression analysis of *Δdcl2* relative to WT. Gene-level differential expression results including log2 fold change, statistical test results, and FDR-adjusted p-values.

**S9 Table.** Pairwise differential expression analysis of *Δpiwi* relative to WT. Gene-level differential expression results including log2 fold change, statistical test results, and FDR-adjusted p-values.

**S1 Fig. Phylogenetic analysis of kinetoplastid Dcl proteins.** The maximum-likelihood phylogenetic tree was performed in IQ-TREE using the PMSF method with 1,000 replicates for ultrafast bootstraps (UFB) and Shimodaira–Hasegawa approximate likelihood ratio test (SH-aLRT). Only SH-aLRT and UFB supports ≥90 and ≥95, respectively, are shown as explained in the graphical legend. Accession numbers are listed in S2 Table.

**S2 Fig. Phylogenetic analysis kinetoplastid Ago1 proteins.** The maximum-likelihood phylogenetic tree was performed in IQ-TREE using the PMSF method with 1,000 replicates for ultrafast bootstraps (UFB) and Shimodaira–Hasegawa approximate likelihood ratio test (SH-aLRT). Only SH-aLRT and UFB supports ≥90 and ≥95, respectively, are shown as explained in the graphical legend. Accession numbers are listed in S2 Table.

**S3 Fig. Phylogenetic analysis of kinetoplastid Rif4 and Rif5 proteins**. The maximum-likelihood phylogenetic tree was performed in IQ-TREE using the PMSF method with 1,000 replicates for ultrafast bootstraps (UFB) and Shimodaira–Hasegawa approximate likelihood ratio test (SH-aLRT). Only SH-aLRT and UFB supports ≥90 and ≥95, respectively, are shown as explained in the graphical legend. Accession numbers are listed inS2 Table.

**S4 Fig. Phylogenetic analysis of kinetoplastid Piwi proteins**. The maximum-likelihood phylogenetic tree was performed in IQ-TREE using the PMSF method with 1,000 replicates for ultrafast bootstraps (UFB) and Shimodaira–Hasegawa approximate likelihood ratio test (SH-aLRT). Only SH-aLRT and UFB supports ≥90 and ≥95, respectively, are shown as explained in the graphical legend. Accession numbers are listed in S2 Table.

**S5 Fig. Validation of target gene KOs via NGS.** Coverage profiles across four KO (*Δago1*, *Δdcl1*, *Δdcl2*, and *Δpiwi*) and WT lines. Each panel shows one target gene with sequencing coverage displayed for all analyzed lines in parallel. The chromosomal attribution is indicated on the left and the chromosomal position below each panel. The deletion sizes for each gene are shown in rectangles.

**S6 Fig. Growth kinetics and morphometry.** *(A)* Growth curves of WT and 4 derived KO lines. Curves were constructed from 3 biological replicates. *(B)* Violin plots of 6 measurements taken from 200 parasites per line. Statistical analyses (multiple *t*-tests) were done in GraphPad Prism (asterisks denote *p*-values as follows: *, < 0.05; **, < 0.01; ***, < 0.001; ns, not significant).

**S7 Fig. Size distribution and nt composition of total small RNA libraries from L. guyanensis RNAi effector KOs.** Stacked bar charts showing 5’ terminal nt composition (left panel) and 3’ terminal nt composition (right panel) across read lengths 15-30 nt for WT, *Δago1*, *Δdcl1*, *Δdcl2*, and *Δpiwi* KO libraries. Colors indicate nucleotide identity. Data represent mean ± SD from WT controls (n = 4) and KO lines generated from two independent clones with four biological replicates each (n = 8 per KO).

**S8 Fig. Quantitative analysis of total vsiRNA abundance across RNAi effector KOs.** Total small RNA reads (21-23 nt) mapping to the LRV1-4 genome (both forward and reverse) were quantified as reads per million (RPM) across WT and KO lines: *Δdcl1, Δdcl2, Δago1, Δpiwi*. Each point represents biological replicates from WT controls (n = 4) and KO lines generated from two independent clones with four biological replicates each (n = 8 per KO). Statistical significance was determined by ordinary one-way ANOVA with uncorrected Fisher’s LSD test comparing each KO to WT control. ****, *p* < 0.0001; *, *p* < 0.05; ns, not significant.

**S9 Fig. Terminal dinucleotide composition of non-template additions in LRV1-4-derived vsiRNAs.** Heatmap showing 3’ terminal dinucleotide addition patterns versus read length (15-30 nt) for soft-clipped reads from WT *L. guyanensis* libraries. Data represent mean counts from WT controls (n = 4) and KO lines generated from two independent clones with four biological replicates each (n = 8 per KO). X-axis shows read length in nucleotides; Y-axis shows all possible dinucleotide combinations.

**S10 Fig. Overlap and fold change relationship of differentially expressed genes across RNAi pathway effector KOs.** *(A)* Venn diagrams showing the overlap of significantly upregulated and downregulated genes in *Δago1*, *Δdcl1*, *Δdcl2*, and *Δpiwi* relative to WT. *(B)* Scatter plot comparing log_2_ fold change values for genes in *Δago1* vs WT and *Δdcl1* vs WT. Points are colored according to significance category: significant only in *Δago1*, significant only in *Δdcl1*, or significant in both comparisons. For readability, labels are limited to representative high-confidence DEGs and are shown by locus suffix; complete pairwise differential expression results are provided in S6-S7 Tables. Dashed lines indicate log_2_ fold change thresholds, and the solid line indicates the linear regression fit with corresponding R^2^ value.

## Notes

### Competing Interest Statement

The authors have declared no competing interest.

