## Supplementary Figure 1 for "Leishmania guyanensis controls endogenous viral replication by a canonical RNA interference pathway"

**Supplementary Figure S1. Phylogenetic analysis of kinetoplastid Dcl proteins.** The maximum-likelihood phylogenetic tree was performed in IQ-TREE using the PMSF method with 1,000 replicates for ultrafast bootstraps (UFB) and Shimodaira–Hasegawa approximate likelihood ratio test (SH-aLRT). Only SH-aLRT and UFB supports ≥90 and ≥95, respectively, are shown as explained in the graphical legend. Accession numbers are listed in Supplementary Table S2.

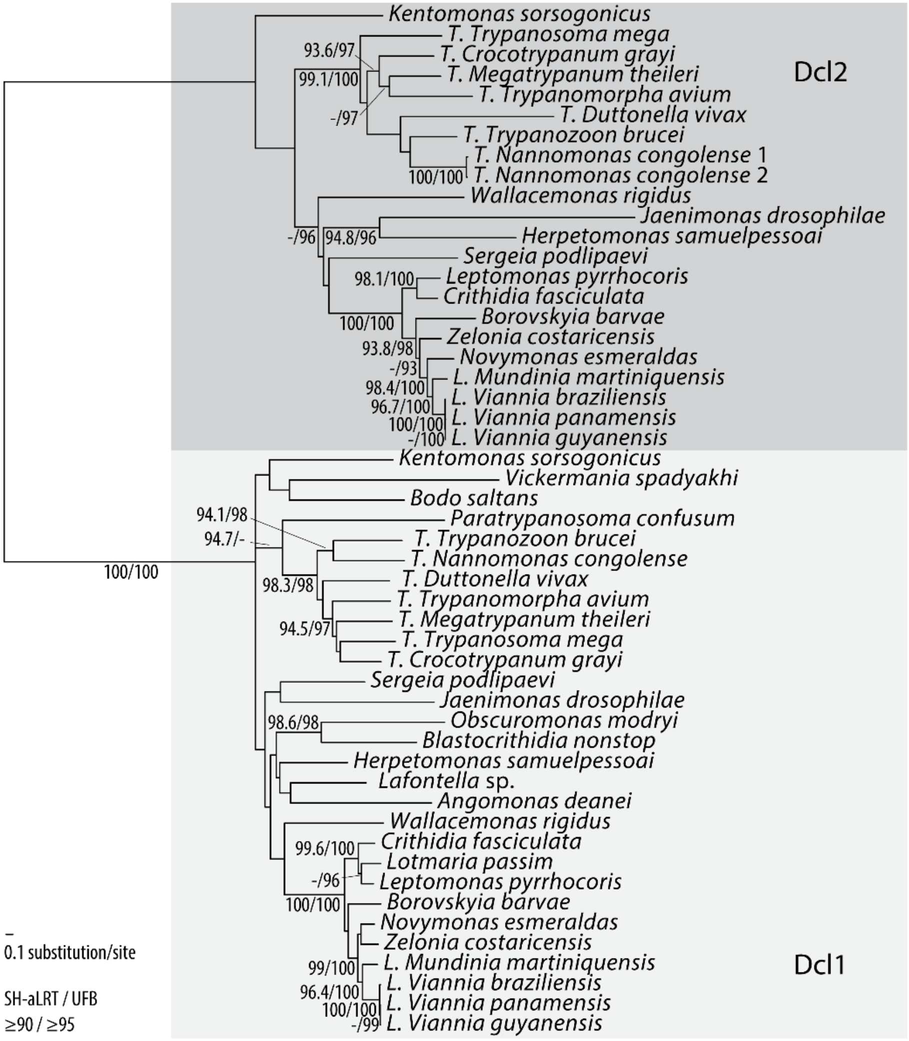
