## Supplementary figures and images for "Leishmania guyanensis controls endogenous viral replication by a canonical RNA interference pathway"

### Supplementary Figure 5

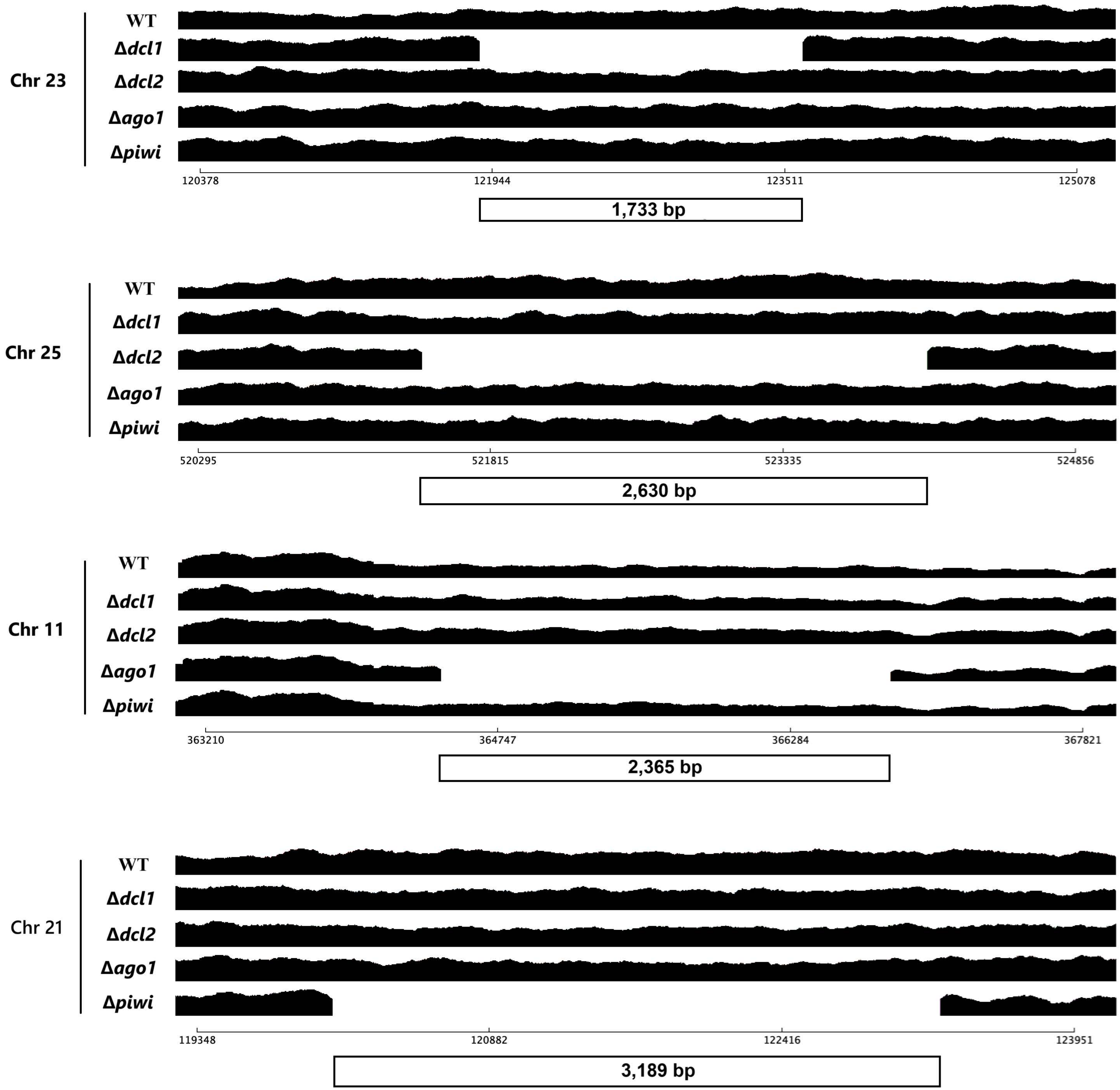
