## Supplementary Figure 6 for "Leishmania guyanensis controls endogenous viral replication by a canonical RNA interference pathway"

**Supplementary Figure S6. Growth kinetics and morphometry. (A)** Growth curves of WT and 4 derived knockout lines. Curves were constructed from 3 biological replicates. **(B)** Violin plots of 6 measurements taken from 200 parasites per line. Statistical analyses (multiple t-tests) were done in GraphPad Prism (asterisks denote p-values as follows: *, < 0.05; **, < 0.01; ***, < 0.001).

**(A)**

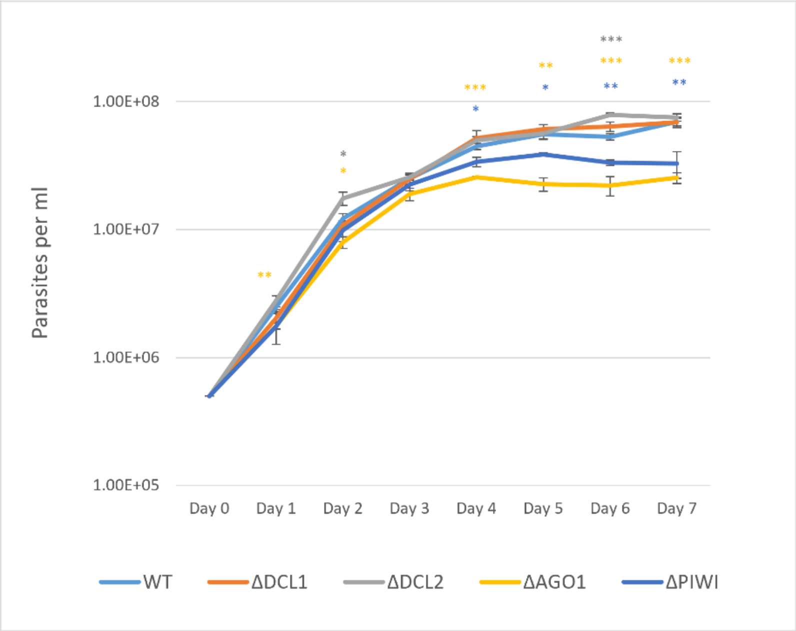


**(B)**

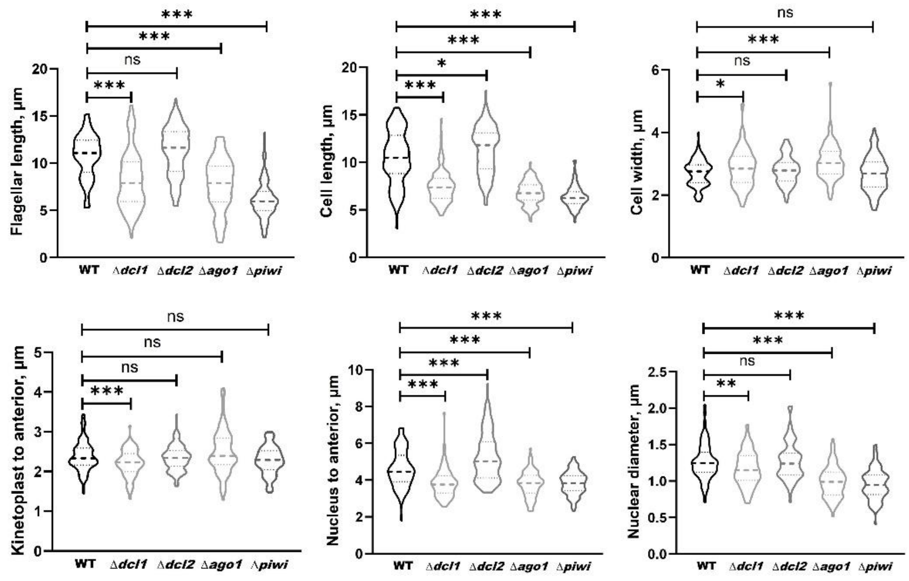
