## Supplementary Figure 7 for "Leishmania guyanensis controls endogenous viral replication by a canonical RNA interference pathway"

**Supplementary Figure S7. Size distribution and nt composition of total small RNA libraries from *L. guyanensis* RNAi effector knockouts**. Stacked bar charts showing 5' terminal nt composition (left panel) and 3' terminal nt composition (right panel) across read lengths 15-30 nt for parental WT, *Δago1*, *Δdcl1*, *Δdcl2*, *Δpiwi* knockout libraries. Colors indicate nucleotide identity: A (orange), C (green), G (yellow), U (blue). Data represent mean ± SD from WT controls (n=4) and knockout lines generated from two independent clones with four biological replicates each (n=8 per knockout).

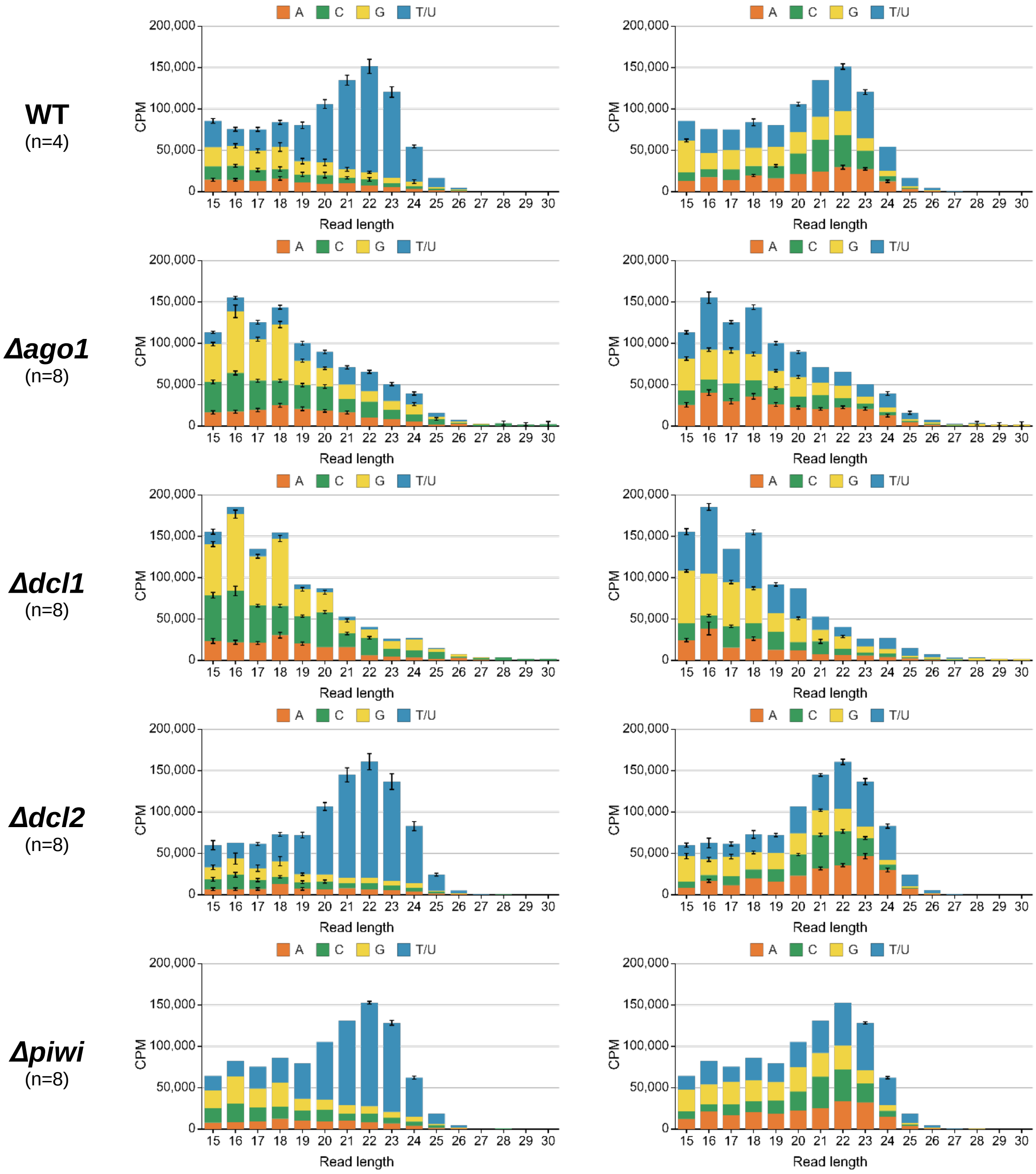
