## Supplementary Figure 8 for "Leishmania guyanensis controls endogenous viral replication by a canonical RNA interference pathway"

**Supplementary Figure S8. Quantitative analysis of total vsiRNA abundance across RNAi effector knockouts.** Total small RNA reads (21-23 nt) mapping to the LRV1-4 genome (both forward and reverse) were quantified as reads per million (RPM, $log_{10}$ scale) across WT and knockout lines: *Δdcl1, Δdcl2, Δago1, Δpiwi*. Each point represents biological replicates from WT controls (n = 4) and knockout lines generated from two independent clones with four biological replicates each (n = 8 per knockout). Statistical significance was determined by ordinary one-way ANOVA with uncorrected Fisher's LSD test comparing each KO to WT control. ****$p$ < 0.0001, *$p$ < 0.05, ns = not significant.

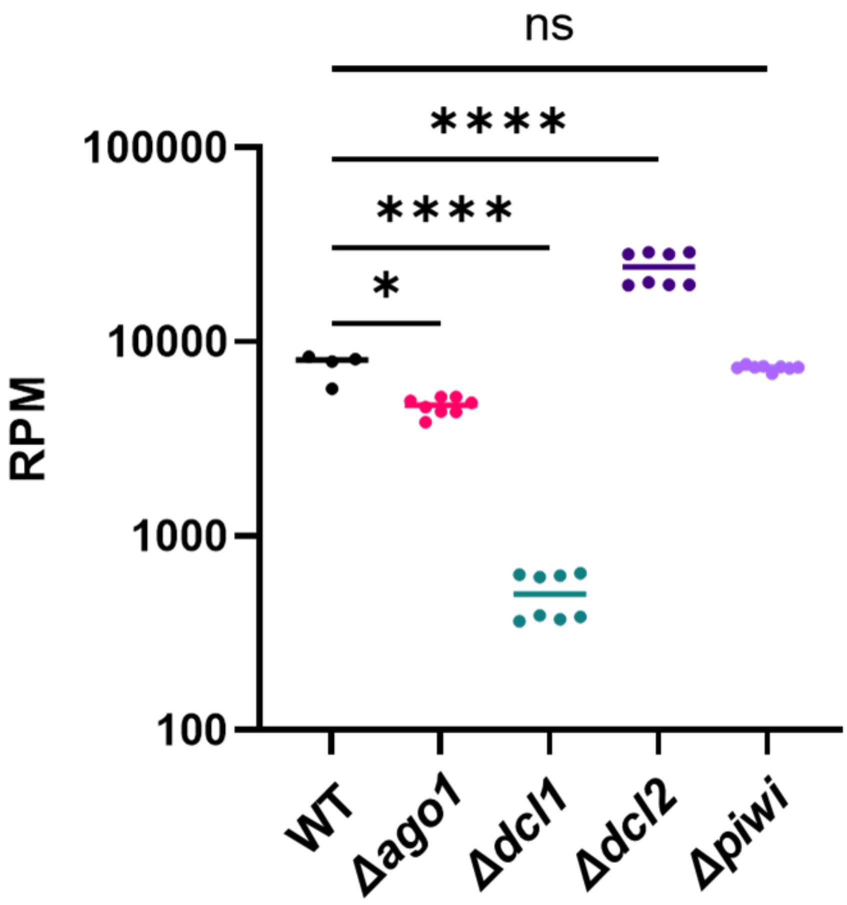
