## Supplementary Figure 9 for "Leishmania guyanensis controls endogenous viral replication by a canonical RNA interference pathway"

**Supplementary Figure S9. Terminal dinucleotide composition of non-template additions in LRV1-4-derived siRNAs.** Heatmap showing 3' terminal dinucleotide addition patterns versus read length (15-30 nt) for soft-clipped reads from *L. guyanensis* libraries. Data represent mean counts per million from WT controls (n = 4) and knockout lines generated from two independent clones with four biological replicates (n = 8 per knockout). Y-axis shows all possible dinucleotide combinations; X-axis shows read length in nucleotides.

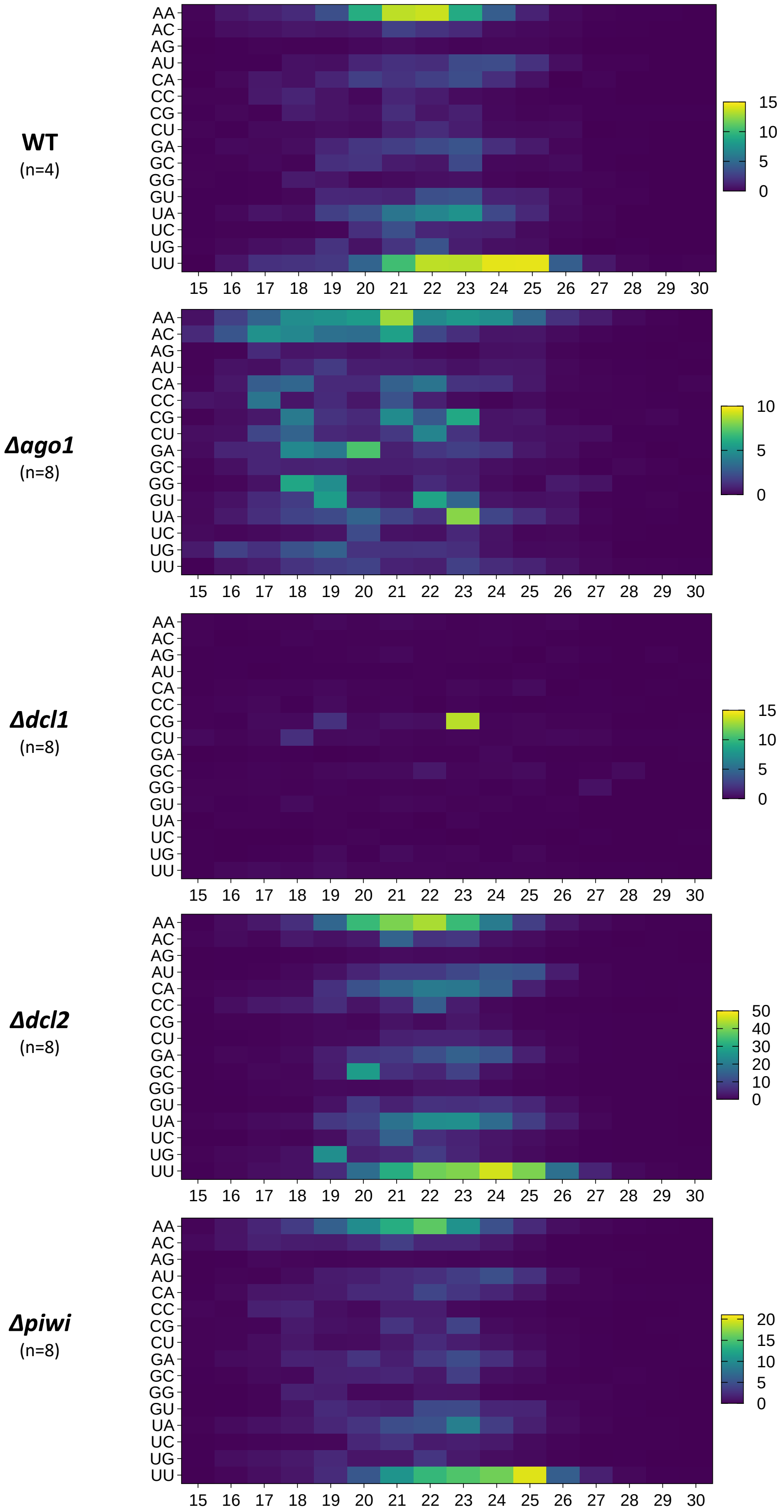
