## Supplementary Figure 10 for "Leishmania guyanensis controls endogenous viral replication by a canonical RNA interference pathway"

**Supplementary Figure S10. Overlap and fold-change relationship of differentially expressed genes across RNAi pathway effector knockouts. (A)** Venn diagrams showing the overlap of significantly upregulated and downregulated genes in *Δago1*, *Δdcl1*, *Δdcl2*, and *Δpiwi* relative to WT. **(B)** Scatter plot comparing $\log_2$ fold-change values for genes in *Δago1* versus WT and *Δdcl1* versus WT. Points are coloured according to significance category: significant only in *Δago1*, significant only in *Δdcl1*, or significant in both comparisons. Gene identifiers are shown for selected genes. Dashed lines indicate $\log_2$ fold-change thresholds, and the solid line indicates the linear regression fit with corresponding $R^2$ value.

A

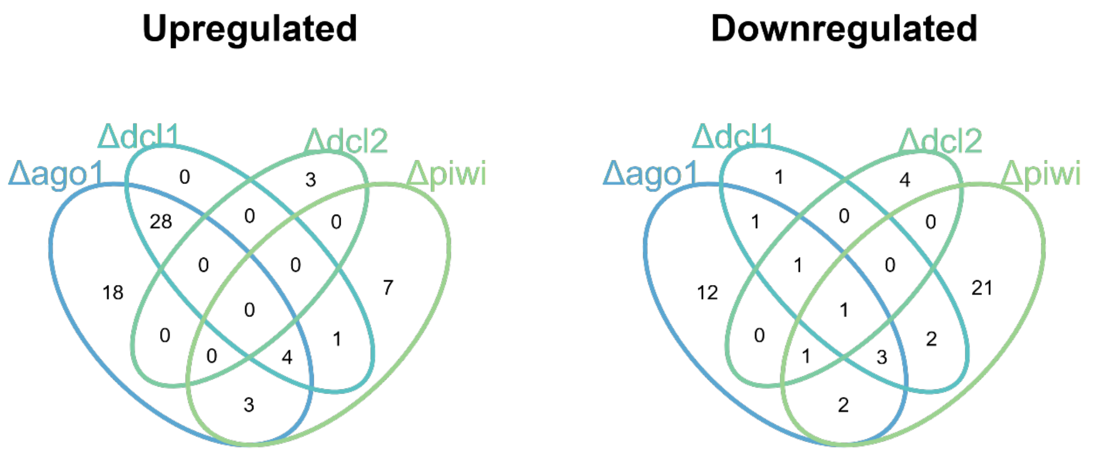


B

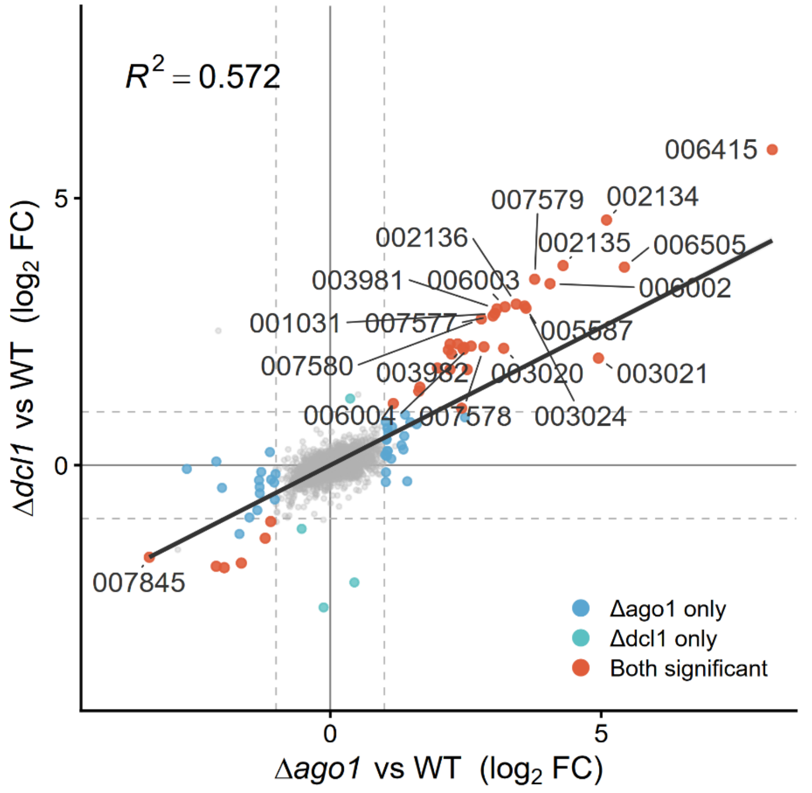
